## Supplementary figures and images for "The evolutionary history of the ancient weevil family Belidae (Coleoptera: Curculionoidea) reveals the marks of Gondwana breakup and major floristic turnovers, including the rise of angiosperms"

### Supplemental Figure 1

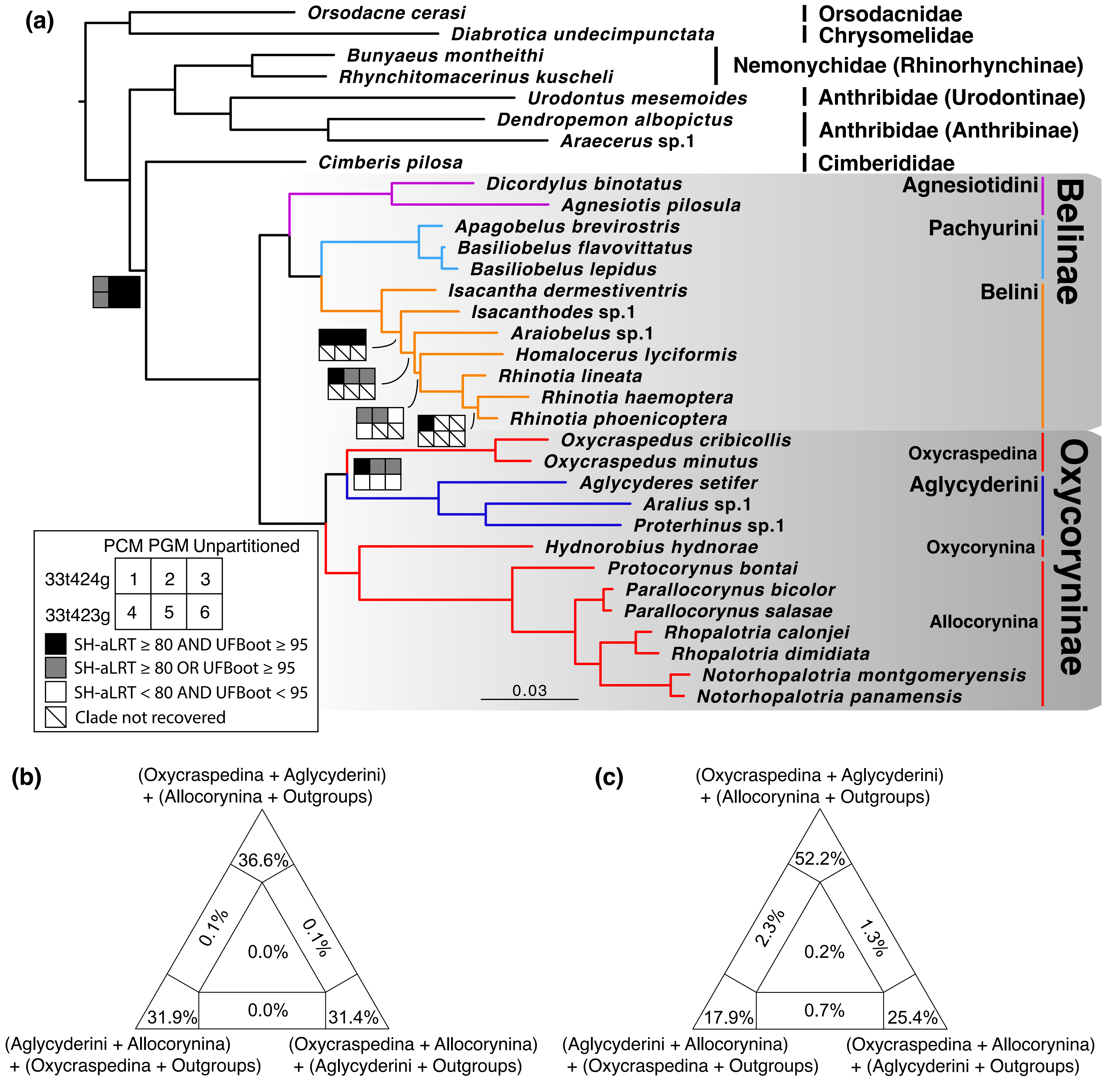

### Supplemental Figure 2

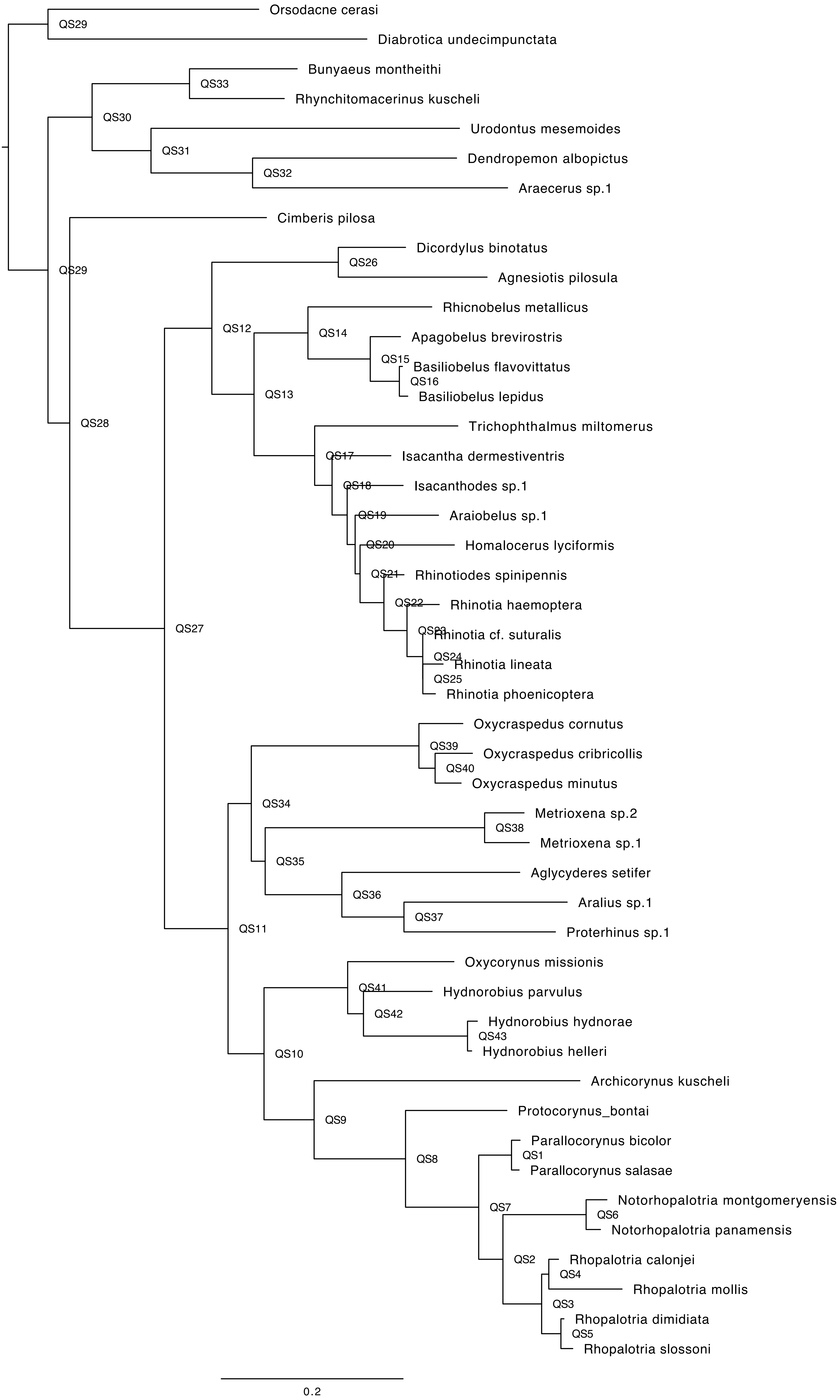
